# Implementing considered elements of standardisation for Time Kill Curve experiments across multiple sites: A European collaboration perspective

**DOI:** 10.64898/2026.06.16.732594

**Authors:** Marie Attwood, Mark Brönstrup, Shampa Das, Hazel L. S. Fuchs, Pippa Griffin, Bettina Hinklemann, Lewis Hoare, Julien Lebrat, Sandrine Marchand, Derry K Mercer, Florian Michel, Alan Noel, Alina Nussbaumer-Proll, Markus Zeitlinger, Alasdair MacGowan

**Affiliations:** North Bristol NHS trust, Bristol, United Kingdom; Helmholtz Centre for Infection Research, Braunschweig, Germany; The University of Liverpool, Liverpool, United Kingdom; Universite De Poitiers, Poitiers; Bioaster Fondation De Cooperation Scientifique, Lyon, France; Medizinische Universitaet Wien, Vienna, Austria

## Abstract

**Background:** The main advantages of Time Kill Curves (TKCs) in antimicrobial drug development are the ability to track bacterial kill and regrowth over time and with varying drug concentrations. Whilst there are guideline documents in place, such as M26-A in CLSI, there remains scope for individual laboratory differences in practice. Here we evaluated several factors which potentially influenced data generated in TKCs.

**Methods:** Firstly, *E. coli* ATCC 25922 was used to determine optimum sampling volume, culture vessel volume, CFU enumeration variance factors and static versus agitated cultures in a single laboratory. Secondly, a ring test comprising of TKCs was performed by six laboratories focusing on: standardised inoculum, static culture and two culture vessel sizes 10 mL and 200 µL. Data analysis was performed to determine consistency within centres and between them.

**Results:** Consistently accurate inocula could be achieved by use of: larger sampling volumes between 100 µL > 20 mL; larger culture vessels volumes (10 mL > 100 µL) and higher inocula (10 ^8^ > 1.5×10 ^5^ CFU). Culture agitation during the TKC experiment resulted in reduced killing compared to static cultures. Reproducibility of TKCs was best between centres when they were performed in 10 mL culture vessels. There was more variability per site when performing TKC in 96 well trays.

**Conclusions:** Technical factors such as preparation of inocula, agitation, vessel size and enumeration of cultures are important variables in performing TKCs that need to be standardised in drug development programmes involving multiple laboratory centres.

## Introduction

With the rise in the burden of antimicrobial resistance (AMR), there has been an increased number of global initiatives launched to tackle this silent pandemic. Interventions, improved strategies and national action plans are all essential elements for the reduction of AMR incidence. However, as a scientific and technical basis, it is imperative to have robust laboratory methodologies to determine efficacy and therapeutic suitability of new antimicrobials.

The GNA NOW project, funded as part of the IHI Innovative Health Initiative, under the global umbrella of IMI’s AMR Accelerator, aims to address the urgent need for antibiotics to treat Gram-negative infections (https://amr-accelerator.eu/project/gna-now/)^1^. This type of collaboration requires cross-centre experimentation, often with simultaneous experimental procedures performed in parallel. Whilst *in vivo* Time Kill Curves (TKCs) are considered gold standard tests for pharmacodynamic (PD) evaluations, there are no published reports of *in vitro* standardisation. There is also no widely accepted information describing data reproducibility for same day (SD) versus different day (DD) replications. The GNA NOW consortium, comprising six independent microbiology laboratories, has previously performed TKC experiments to establish if SD replication and DD replications are consistent^2^ . It also tested if same centre (intra-centre) and cross site (inter-centre) replications were consistent. We found that intra-centre replications (SD and DD) were statistically consistent with each other, whereas inter-centre replications were not. It was hypothesized that inter-centre variation could be due to differences in methodology, specifically inoculum preparation, culture vessel volume and culture conditions.

In this study, we aimed to establish if standardisation of TKC methodology will produce more robust, reproducible and reliable data. First, a single site performed comparison experiments on the following: the impact of sampling volume size and of culture vessel volume size on reproducibility and the impact of culture conditions (static vs agitated) on bacterial killing.

Results from these studies informed decisions on elements that should be standardised. Subsequently, the six independent microbiology laboratories performed TKC experiments under identical conditions; the results of these efforts are reported below.

## Material and Methods

### Single site – replicates n = 6

#### Bacterial isolate, antimicrobial and media

We tested *E. coli* ATCC 25922 (meropenem MIC = 0.03 mg/L) with the multiple variables being evaluated. We sourced meropenem from ACS Dobfar (1g powder for solution for injection/infusion), the solvent and diluent were used as stated by the manufacturer (H_2_O, and potency was 100%). Targets were set as multiples of MIC; x0 (growth control), x1, x2, x4, x8, x16. The media employed was Muller Hinton Broth II – BD BBL™ Dehydrated culture Media, which was also used for previous evaluation TKC experiments. Certificates of compliance were BS12, CLSI, CMPH2, MCM9 and the manufactures reconstitution instructions were carried out, followed by sterilisation by autoclaving. Environmental conditions stated for TKC culture were aerobic conditions at 37°C with and without agitation. Agitation was set at 150RPM with a throw of 2cm to ensure consistent and reproducible mixing conditions.

#### Sampling volume size and reproducibility

The target inoculum was established at 1.5 × 10^6^ CFU/mL, and initial CFU enumeration sampling was conducted 15 mins after inoculum administration. CFU enumeration sampling size varied between 20 µL, 50 µL and 100 µL for CFU enumeration for a set culture vessel volume. CFU enumeration was then performed using a spiral plater (Don Whitley Scientific, Yorkshire, UK) onto Muller-Hinton II agar after incubation for 20 h. Meropenem was not added to these sets of experiments.

#### Culture vessel volume, inoculum CFU/mL density and reproducibility

Target inocula were set at 1.5 ×10^8^, 1.5 × 10^7^, 1.5 × 10^6^, 1.5 × 10^5^ CFU/mL for all culture vessel volumes; 10 mL, 5 mL, 2 mL, 200 µL and 100 µL. Meropenem was not added to these sets of experiments.

#### Culture vessel volume size, sampling volume size and reproducibility

Target inoculum as stated above, sampling (for CFU enumeration) started 15 mins after inoculum administration. Total volume of culture vessel varied between 100 µL, 200 µL, 2 mL, 5 mL and 10 mL. Sampling volume size was a set volume of 100 µL. Then CFU enumeration was performed with a spiral plater (Don Whitley Scientific, Yorkshire, UK) onto Muller-Hinton II agar after incubation for 20 h. Meropenem was not added to these sets of experiments.

#### Culture conditions and reproducibility

TKC with initial inoculum of 1.5 × 10^6^ CFU/mL using a 10mL static culture vessel. This was compared to an agitated (150 RPM) TKC with initial inoculum of 1.5 × 10^6^ CFU/mL using a 10 mL culture vessel. Meropenem was added to these simulations in MIC multiples of x1, x2, x4, x8, x16.

#### Experimental sites

Listed in alphabetical order; Bioaster Fondation De Cooperation Scientifique, Lyon, France. Helmholtz Centre for Infection Research, Braunschweig, Germany. Medizinische Universitaet Wien, Vienna, Austria. North Bristol NHS trust, Bristol, United Kingdom.

Universite De Poitiers, Poitiers, France. University of Liverpool, Liverpool, United Kingdom.

#### Experimental volumes, conditions and data points tested (Ring test)

The activity of meropenem against the *E. coli* bacterial isolate ATCC 25922 was evaluated using the previously defined MIC multiples. Predefined total volumes of culture vessels were used in these TKC experiments, 10 mL glass universal was compared with 200 µL plastic microtiter trays. All TKC cultures were incubated under static conditions. Timepoints for bacterial count enumeration were at hours 0, 2, 4, 6, 8 and 24. All results were reported in CFU/mL. All simulations were performed in triplicate. Centres which did not meet inoculum criteria (5.5 × 10^5^ CFU/mL – 5.5 × 10^6^ CFU/mL) were excluded.

#### Statistical tests and assumptions

To compare the distribution between experimental inoculum repeats, an analysis of variance the two-way ANOVA statistical test was used. The assumptions are that the data is normally distributed for same day versus different day intra-site replications and inter-site replications. Target bacterial inoculum at timepoint 0 (T0) was set at 1.5×10^6^ CFU/mL to establish baseline CFU/mL and prior to antibiotic exposure. The 1st null hypothesis, H0, is the expectation that repeated measures at T0 intra-site are consistent. The alternative hypothesis, H1, is that the repeated measures at T0 intra-site (across MIC multiplies) are statistically different. The second null hypothesis, H0, is that repeated measures at T0 inter-site are consistent. The alternative second null hypothesis, H1, is that repeated measures at T0 inter-site are statistically different. Normal distribution cannot be assumed once bacteria have been exposure to antibiotic and therefore another statistical test was applied for these evaluations.

Friedman test was employed to determine any statistical difference between intra and inter centre same day and different day data. In this application of the Friedman test, the null hypothesis, H_0_, is the expectation that concentration of antibiotic has no effect on the trend of total number CFU/mL. Therefore, the alternate hypothesis, H_1_, states that antibiotic concentration affects the trend of CFU/mL. To assess the rate of bacterial kill, the analysis focused on T4 and T24 time points. Friedman test values at timepoints previously specified were compared against the threshold value derived from Chi-squared (χ ^2^) distribution tables. If the Friedman test values exceed this threshold, it is indicative that the replications within individual sites and across sites are consistent with each other. To summate this the higher the test statistic which exceeds the threshold the more consistent the data points. The number of experiments using different antibiotic concentrations in each group, k, was 6, so the reported χ^2^ (DoF = 5, a = 0.05) threshold value was Fr_threshold_ = 11.070.

## Results

### Single centre evaluations

The effect of sampling volume size and the total volume of the culture vessel on the reproducibility of a 1.5 × 10^6^ CFU inoculum target is shown in Figure 1. In general, as the sampling volume increased from 20 µl to 100 µL, the variability of the measured bacterial load decreased. In addition, an increase of the total vessel volume from 100 µL to 10 mL led to decreased variability in the measured bacterial density.

**Figure 1.**
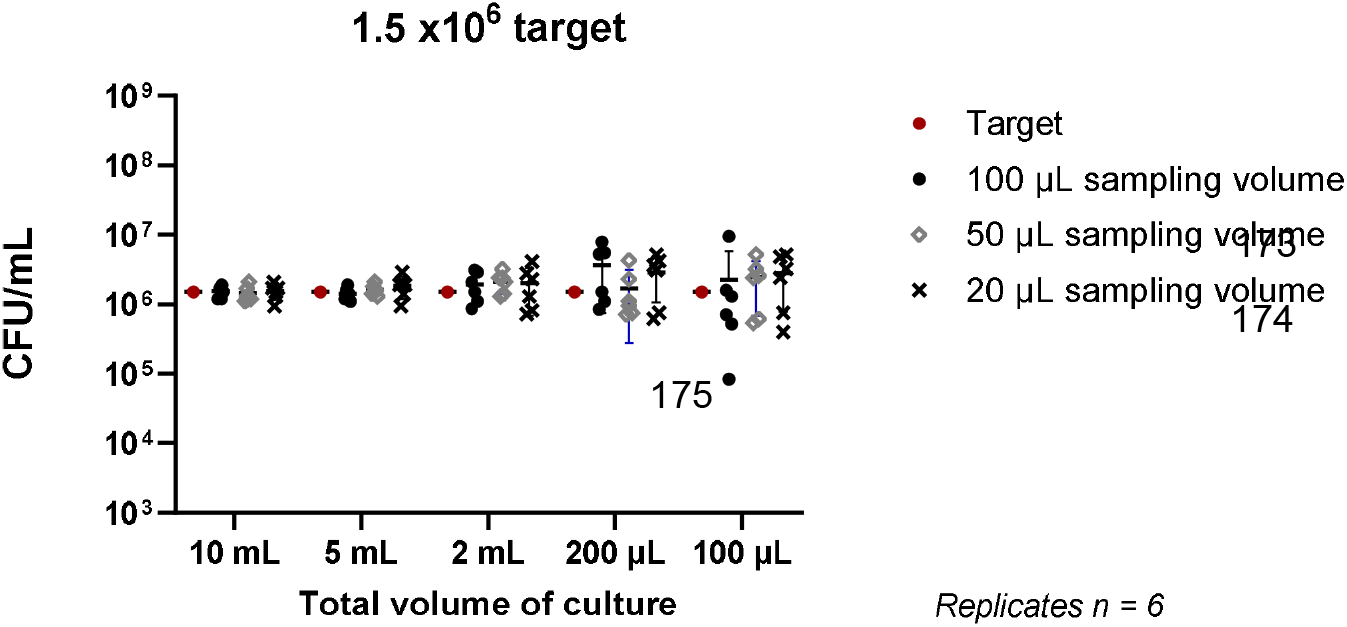
The effect of sampling volume size and total vessel volume on reproducibility.

The impact of bacterial inoculum size and the total volume of the culture vessel on the measured bacterial density is shown on Figure 2. As the total volume of the culture vessel increased from 100 µL to 10 mL, the variability in bacterial counts decreased. This was particularly marked for lower target inocula.

**Figure 2.**
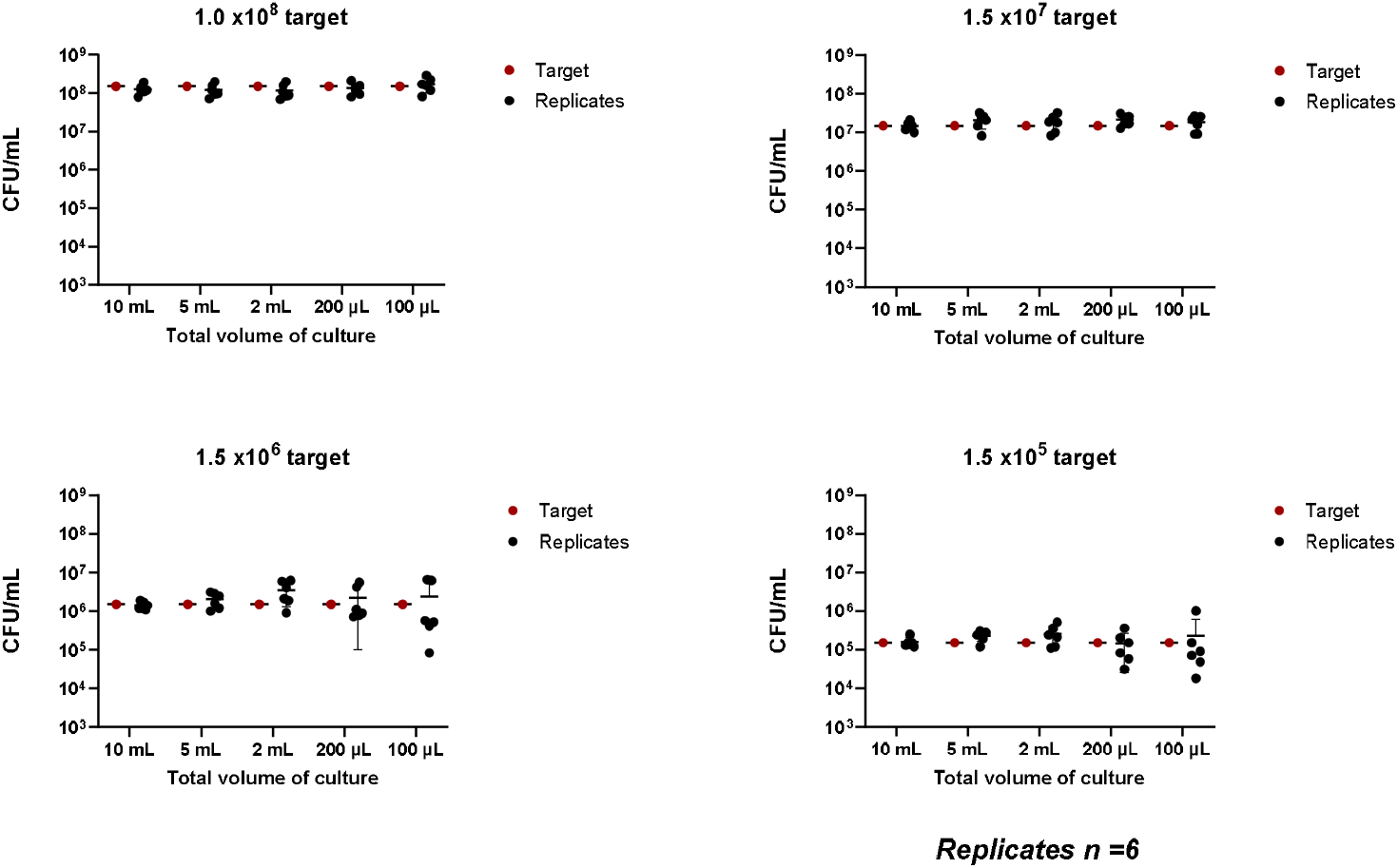
The effect of total vessel volume and target inoculum size on reproducibility.

The impact of agitation of the culture vessel is shown on Figure 3. Agitation was associated with reduced bacterial kill by meropenem at concentrations of x4, x8 and x16 the MIC.

**Figure 3.**
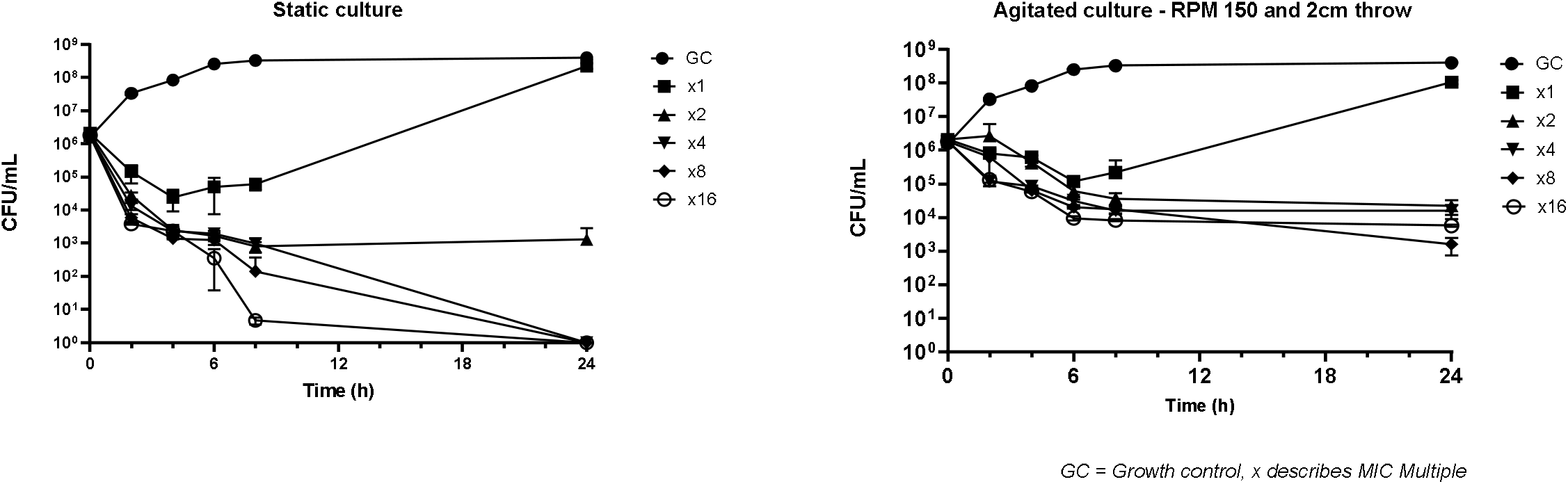
The effect of culture agitation on reproducibility, *E. coli* 25922.

**Figure 4.**
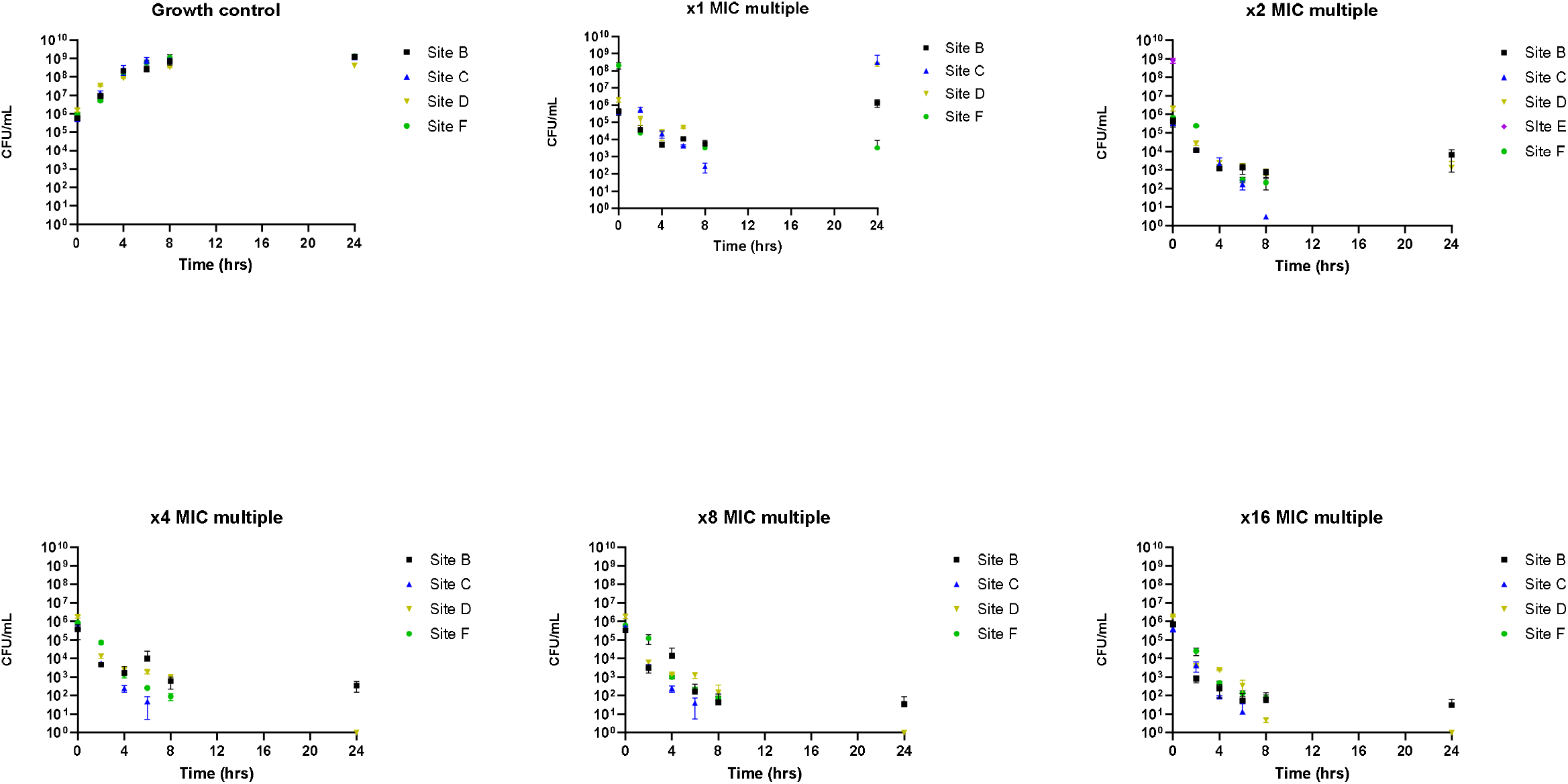
Inter-centre 10 mL same day comparison.

**Figure 5.**
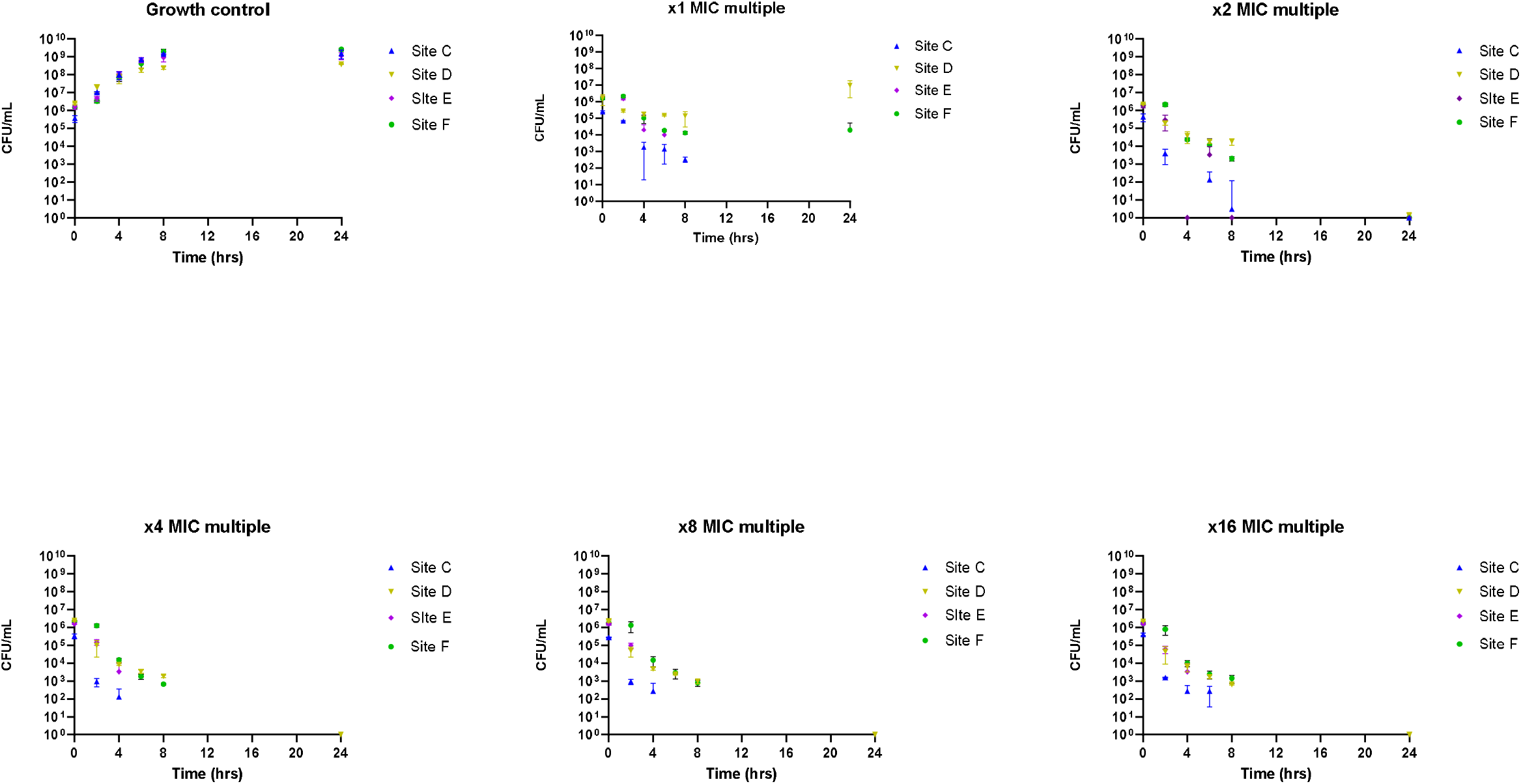
Inter-centre 200 µL same day comparison.

### Multi-centre evaluations

### Multi centre evaluations data analysis

Introducing inoculum limits restricts variability and inevitably increases the reproducibility of same centre and cross centre inocula comparisons. This has been proven as no statistically significant differences between inocula under either testing condition where observed. ANOVA details can be found in supplementary tables 1 and 2.

#### T2+T4 and T24 same day analysis

Friedman test statistics for each laboratory included two recorded timepoints at 2 and 4 hours. These times points were used to assess bacterial kill. Test statistics are equally important at later timepoints, and regrowth measured by 8- and 24-hour data to determine compound efficacy and possible emergence of resistance. All test statistics were critically compared to a threshold value of 11.07 from χ^2^. Dotted line depicts this threshold value.

Intra-site comparison shows that bacterial kill and regrowth trends were consistent for all site replicates when using 10 mL culture vessels for both T2+4 and T24 hours (Figure 6 and 7). Data obtained from 200 µL vessels displayed inconsistencies between sites when assessing kill (Figure 6). A higher level of consistency per site was obtained for the regrowth data at 24 hours (Figure 7).

**Figure 6.**
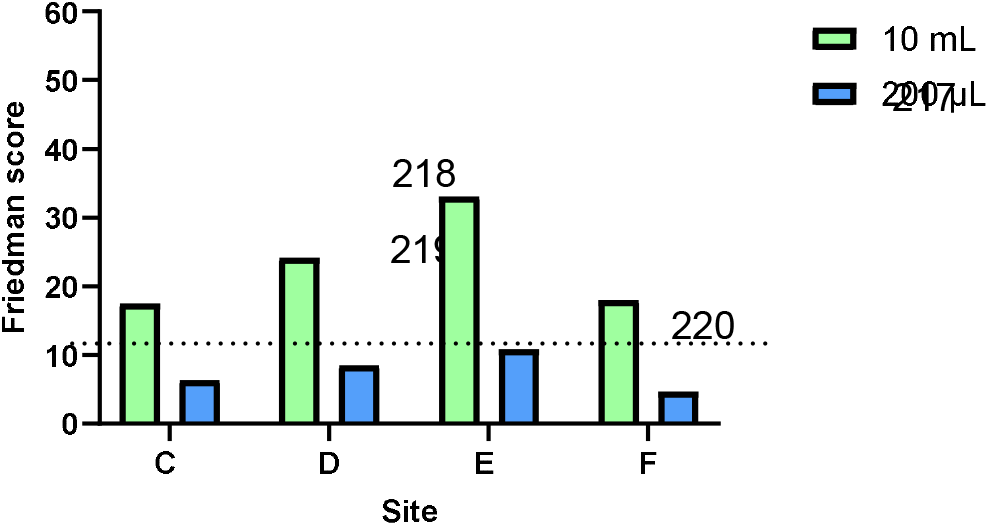
Intra-centre same day comparisons for rapid kill hour 2 and 4.

**Figure 7.**
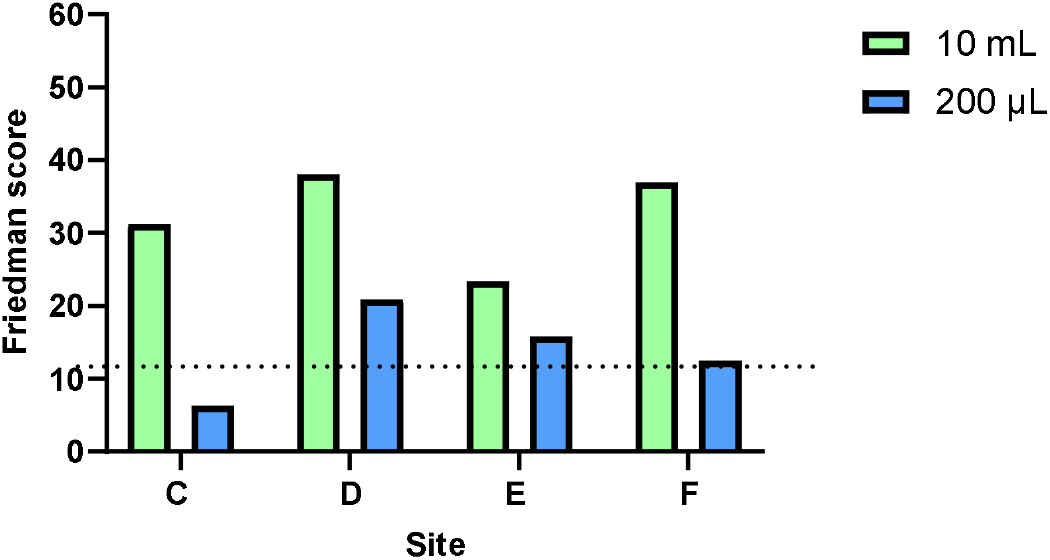
Intra-centre same day comparisons for regrowth hours 8 and 24.

The inter-centre comparison shows that use of a 10 mL culture vessel had more consistent trends across sites in relation to rapid kill and regrowth (Figure 8 and 9). Results obtained from 200 µL total vessel volume showed good consistency across sites at 24 hours but had variability across sites when assessing kill (Figure 8).

**Figure 8.**
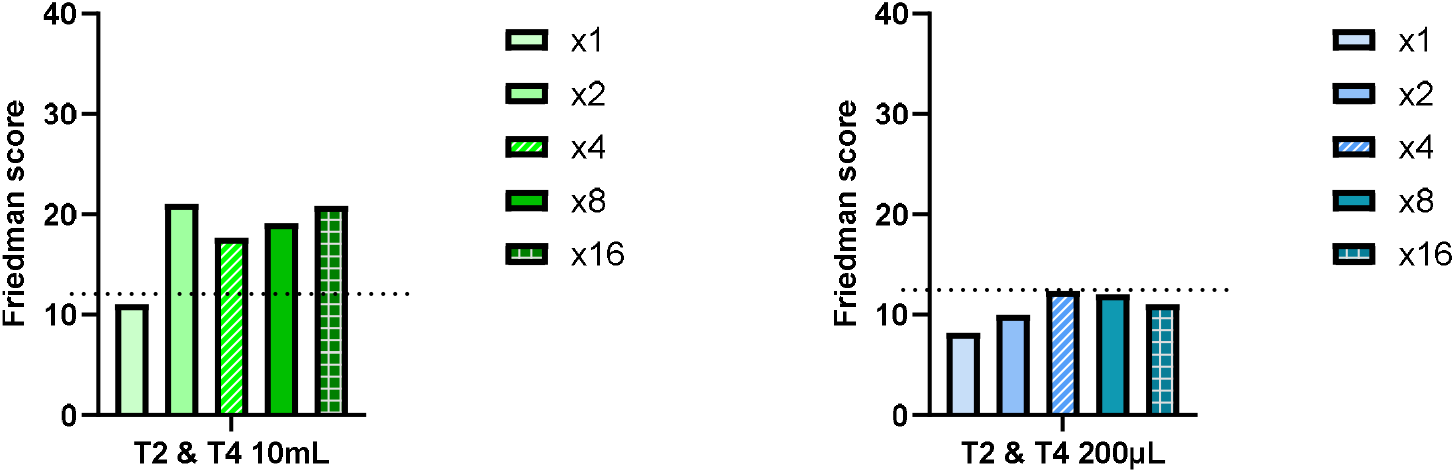
Inter-site same day comparison 10 mL and 200 µL total volume cultures for rapid kill.

**Figure 9.**
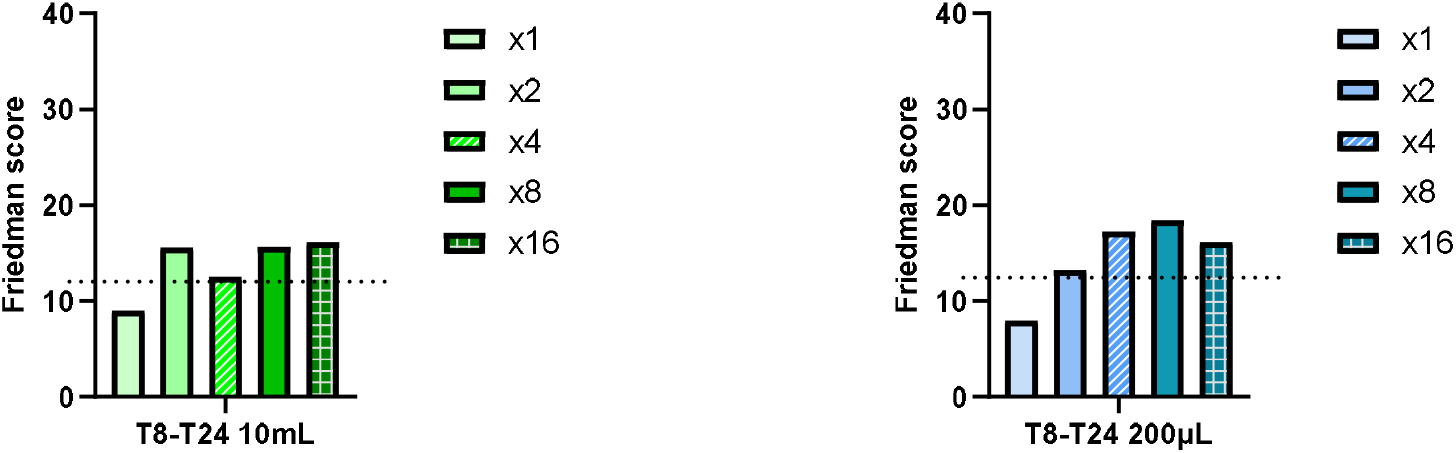
Inter-site same day comparisons 10mL and 200µL total volume cultures for regrowth.

## Discussion

Time kill curves have the advantage that they allow the tracking of bacterial kill and regrowth over time and at different concentrations. Whilst there are guideline documents in place such as M26-A in CLSI^3,4^, there remains scope for individual inter-laboratory variations.

Therefore, the use by individual laboratories of specific equipment and/or techniques may influence the development of TKCs which in turn affect the data generated and interpretation. The largest variation in TKCs generally occurred at lower (x1 – x4) multiples of MIC compared to the extremes of antibiotic concentrations, i.e. the growth control or X8 and x16 MIC multiples, which exhibited similar trends regardless of culture vessel size (Figure 6-9). This variation could be related to ambiguities of minimal inhibitory concentration determination i.e. by doubling dilutions instead of using a linear dilution scale. As previously reported^56^, the inoculum size plays an important factor in determining antibiotic effect. We have addressed this factor and shown that is possible to improve intra/inter centres same day and different day reproducibility by setting inoculum targets with tolerance limits.

TKCs performed in 96 well trays are commonly used in order to run multiple experiments in an efficient manner. However, there has been criticism of this approach due to impaired nutrient availability, reducing the accuracy of results and the translational value in correlations with clinical outcomes^7^. This evaluation has demonstrated similar general kill trends, but with wider variability between data points for both rapid kill and regrowth assessments for small vessel volumes. Specifically, we have determined a larger margin of error when performing TKCs in volumes <200 µL, as pipetting errors in relation to drug dispensing / inoculum are magnified. These smaller volumes of TKC also require smaller sampling volumes, which in turn, increases variability in CFU enumeration associated with pipetting errors. In turn, we have observed that larger TKC culture volumes allow for larger sampling volumes which reduce variability in CFU determination. On the other hand, the use of larger culture vessels increases the amount of compound required for the assay. While larger volume TKC cultures lack dynamically changing antibiotic concentrations, there is translational evidence to support *in vitro* 10 mL TKCs, *in vivo* TKCs and dilutional *in vitro* models correlations^8–10^.

As new classes of antimicrobials, peptides and other novel therapeutics often exhibit specific anti-infective and bacterial interaction characteristics, it is important to remain flexible in altering specific test conditions. For example, the addition of specific serum to the test conditions was indicated in the case of zosurabalpin^11^ to better simulate in vivo conditions. In addition to technical factors discussed above, it is also important to try and develop a consensus on what is the desired outputs in terms of end points should be, for example, fast rate of kill at short time points or minimal regrowth at longer time points.

Previous work^x^ has highlighted the importance of establishing precise experimental protocols upon project commencement and assessment of the factors that can have an influence on quantitative data. In the case of these TKCs, influencing factors are total culture volume size, sampling volume size and static vs agitated cultures. Whilst one solution could be the for formation of a ‘central testing site’, this perhaps would not be appropriate for all collaboration projects. This study also presents a solution for determination of TKCs across multiple sites. It supports the feasibility of multi-centre experiments by establishing a strong foundation through well-considered, bespoke and defined experimental protocols.

## Acknowledgement

The GNA NOW consortium has received funding from the Innovative Medicines Initiative 2 Joint Undertaking under Grant Agreement n°853979. This Joint Undertaking receives the support from the European Union’s Horizon 2020 research and innovation programme and EFPIA.

## Disclaimer

Funded by the European Union, the private members, and those contributing partners of the IMI JU. Views and opinions expressed are however those of the author(s) only and do not necessarily reflect those of the aforementioned parties. Neither of the aforementioned parties can be held responsible for them.

## Transparency Declarations

MA, PG, ARN and APM holds research grants/activities with Merck, Shionogi, InfectoPharm, GSK, Roche, BioVersys, VenatoRx Pharmaceuticals, iFAST, Technomede, Oxford Drug Design, JPIAMR and NIHR; APM provides consultancy advice to Shionogi, Roche, Bicycle Therapeutics and BioVersys. DM provides consultancy advice to INCATE, Selmod GmbH and University of Basilicata. All other authors declare no competing interests

## References

1. ihi.europa.eu/project-results/project-factsheets/gna-now.

2. Attwood, M. et al. Is there a need to implement standardisation into in vitro antimicrobial evaluation systems? A European collaboration perspective. Preprint at 10.64898/2026.06.11.731574 (2026).

3. https://clsi.org/shop/standards/m26.

4. https://standards.iteh.ai.

5. Mould, F. L., Kliem, K. E., Morgan, R. & Mauricio, R. M. In vitro microbial inoculum: A review of its function and properties. Anim. Feed Sci. Technol. 123–124, 31–50 (2005).

6. Gloede, J., Scheerans, C., Derendorf, H. & Kloft, C. In vitro pharmacodynamic models to determine the effect of antibacterial drugs. Journal of Antimicrobial Chemotherapy 65, 186–201 (2010).

7. Mueller, M., De La Peña, A. & Derendorf, H. Issues in Pharmacokinetics and Pharmacodynamics of Anti-Infective Agents: Kill Curves versus MIC. Antimicrobial Agents and Chemotherapy vol. 48 369–377 Preprint at 10.1128/AAC.48.2.369-377.2004 (2004).

8. MacGowan, A. P. et al. The pharmacodynamics of fosfomycin in combination with meropenem against Klebsiella pneumoniae studied in an in vitro model of infection. Journal of Antimicrobial Chemotherapy 80, 967–975 (2025).

9. Noel, A. R., Attwood, M., Bowker, K. E. & MacGowan, A. P. Pharmacodynamics of taniborbactam in combination with cefepime studied in an in vitro model of infection. Int. J. Antimicrob. Agents 64, 107304 (2024).

10. Muller, A. E. et al. Cefepime pharmacodynamic targets against Enterobacterales employing neutropenic murine lung infection and in vitro pharmacokinetic models. Journal of Antimicrobial Chemotherapy 77, 3504–3509 (2022).

11. www.clinicaltrialsarena.com/analyst-comment/zosurabalpin-promise-antibacterial-resistance/?cf-view.

