## Supplementary material for "Implementing considered elements of standardisation for Time Kill Curve experiments across multiple sites: A European collaboration perspective": sup material

Supplementary materials

**Table S1 - *E. coli* 25922 10mL culture vessel T0**

| *E. coli* 25922 - 10mL | | | | | | |
| --- | --- | --- | --- | --- | --- | --- |
| **ANOVA table** | **SS** | **DF** | **MS** | **F (DFn, DFd)** | **P value** | **Significant** |
| Treatment (between columns) | 4.70E+17 | 3 | 1.57E+17 | F (3, 20) = 0.7295 | P=0.5464 | No |
| Residual (within columns) | 4.29E+18 | 20 | 2.15E+17 |  |  |  |
| Total | 4.76E+18 | 23 |  |  |  |  |

**Table S2 - *E. coli* 25922 200µL 96 well tray culture vessel T0**

| *E. coli* 25922 - 200µL | | | | | | |
| --- | --- | --- | --- | --- | --- | --- |
| **ANOVA table** | **SS** | **DF** | **MS** | **F (DFn, DFd)** | **P value** | **Significant** |
| Treatment (between columns) | 1.54E+18 | 3 | 5.14E+17 | F (3, 20) = 0.9698 | P=0.4265 | No |
| Residual (within columns) | 1.06E+19 | 20 | 5.30E+17 |  |  |  |
| Total | 1.22E+19 | 23 |  |  |  |  |

SS – sum of squares

DF- degrees of freedom

MS – mean of squares

F(DFN, DFd) – F ratio
